# Sturgeons: An Evolutionary Insight to The α-globin Protein Speciation and Diversification

**DOI:** 10.1101/2022.06.11.495750

**Authors:** Shohreh Ariaeenejad, Kaveh Kavousi, Elaheh Elahi, Ali Mohammad Banaei-Moghaddam, Ali A. Moosavi-Movahedi

## Abstract

Sturgeons are living fossils and among the oldest surviving vertebrate species, and Caspian Sea is habitat of the vast majority of sturgeon species. It is tempting to assume that speciation of sturgeons occurred in response to the severe environmental changes during the formation of Caspian Sea from the ancient Paratethys Sea. Hemoglobin (Hb) reflects the evolutionary history of fishes. Changes in the Hb sequences will affect the protein structure and consequently the vital function of oxygen delivery. As complete sequence data for any sturgeon hemoglobin gene or protein was hitherto not available, we determined the complete gene and protein sequences of an α-globin chain (as one of the constituents of Hb complex) from six Caspian Sea sturgeons of different species and compared these sequences mutually and with all available fish α-globin sequences. The phylogenetic analyses based on alpha globin sequences can highlight the essentiality of proper changes in globin chains in the speciation process. Estimating the divergence time of α-globin genes for different fish families discloses important facts about the rate of evolution and decodes. The results support the hypothesis of simultaneity between the time of morphological speciation of fish families and their α-globin divergence. The correlation and concurrency between morphological and molecular changes can be interpreted as a sign of struggling to acquire more fitness to changing environment.

## Background

Sturgeons (family *Acipenseridae*) constitute a fish family within the order Acipenseriformes. The order Acipenseriformes belongs to the class Actinopterygii (ray-finned fishes). The study of their Hbs is of interest for two reasons. First, the evolutionary history of Acipenseriformes goes back to the Triassic period and second, they are among the most ancient of actinopterygians. Sturgeons, in evolution fit somewhere between sharks and bony fishes. They share a largely cartilaginous endoskeleton and similar jaw suspension with sharks. Contrary, although they are not ancestral to modern bony fishes and their skeleton is almost wholly cartilaginous, they have a series of external bony plates, called scutes along their backs and sides [1–3]. These living fossils are among the oldest surviving vertebrates [4].The sturgeon species have been described based on morphological characters. All of them are living in the Northern Hemisphere. Most sturgeons are anadromous bottom-feeders, spawning upstream and feeding in river deltas and estuaries [5]. Therefore, compared to the fishes exclusively living in the ocean, sturgeons acquired more adaptation for living under different condition of marine and fresh water. Worldwide, there are 25 sturgeon species in four different genera, among them 17 Acipenser species and two Huso species [3,6,7]. Six of these including *Acipenser persicus, A.gueldenstaedtii, A.ruthenus, A.nudiventris, A.stellatus*, and *Huso huso* are found in the Caspian Sea. All species except *A.ruthenusoccur* throughout the Caspian Sea, but mostly feed and spend the winter in the southern areas of the basin. The biggest spawning population also concentrates in the rivers including Kura, Ural, Volga, Atrak, Samur, Sefid-Rud, Gorgan-Rud, and Tajan.

*A.persicus* is endemic to the Caspian basin, most abundant in the southern part [8]. *A. nudiventris* mainly inhabits the Caspian and the Black Sea [9]. *A.stellatus, H.huso*, and *A. gueldenstaedtii*, besides the Caspian and Black Sea, inhabit the Sea of Azov [5]. *A.ruthenus* is a freshwater species inhabiting the rivers flowing into the Caspian, Black, Baltic, White, Barents, Kara seas and the Sea of Azov [9].

*A.persicus* (Persian sturgeon) and *A.gueldenstaedtii* (Russian sturgeon) were earlier classified as a single species [10] because these sturgeons are morphologically very similar [9].

Among vertebrates, fishes are particularly suitable for studying adaptive processes because of their evolutionary position and the variability regarding their environmental conditions. At the molecular level, adaptations are based on changes of nucleotide sequences of genes and consequently of amino acid sequences of encoded proteins [11,12]. The changes in amino acid sequences affect protein structure and influence protein function. Historically, Hb was frequently studied for elucidation of structure-function relationships [13–16]. Hb is expected to be a preferred target of adaptive processes in fish. Its function in delivering oxygen to tissues for aerobic metabolism is essential for their survival and this function is affected by the enormously variable conditions in which fishes live [17–21]. The conditions include temperature, pressure, salinity, and oxygen availability. The extent of variability in fish Hbs is a footprint of evolutionary and adaptive processes that have affected these molecules [22,23].

The importance of globin in molecular evolutionary clock and timetree studies goes back to the seminal paper of Zuckerkandl and Pauling [24]. In 1962, they estimated the divergence time of four globin chains based on the molecular clock calibrated using the number of observed differences between human and horse α-globin proteins and the divergence time between two species based on fossil records. Scientists had observed that the number of amino acid differences between some important proteins (globin, cytochrome c and fibrinopeptides) from different species is correlated with their divergence time based on the fossil records [25].

It is widely accepted that the sensitivity of a protein in estimating evolutionary time would increase with time of evolutionary divergence of the species being analyzed, protein sequence length, and the substitution rate of the protein [26]. Although the accuracy of molecular clocks would be improved by using various proteins of different types, in this study, due to the lack of an existing sequence of other proper proteins for all investigated fish species, we studied only α-globin.

Various studies have been focused on the globin chains for determining the evolutionary position of fish species [27–30]. But, to the best of our knowledge, this is the first time that the globin chains of sturgeons are used for this purpose.

One of the main goals of this study was to perceive the impact of environmental changes on the rate of molecular clock ticks. For this purpose, we need a protein with important function and proven adaptability that is sensitive to environmental conditions. Globin, with such properties, was an appropriate candidate. Between two main clusters of globin genes, namely α-like and β-like gene families, the consistent location of α-like globin genes in the *MN* locus (the locus flanked on one side by the genes *MPG* and *NPRL3*) in gnathostomes indicates a more stable history than that of the β-like globin genes. This greater stability is also seen in the composition and expression patterns of the α-like globin genes [31]. Therefore, the α-globin was chosen to test the hypothesis that the molecular clock rate for highly sensitive functional proteins can be variable during evolution in response to the changing environment.

Considering that some species such as A.*persicus* are endemic of Caspian Sea, we were interested in analyzing their α-globin sequences to investigate the evolutionary history after recent geological events leading to the formation of the Caspian Sea. So far, phylogenetic trees based on available α-globin sequences and morphological features were not investigated together in previous studies of sturgeons. Complete sequence data on any single sturgeon Hb gene or protein was hitherto not available. Therefore here, we provide complete gene sequences of the α-globin chain of six Caspian Sea sturgeons and amino acid sequences derived in silico. The GenBank accession numbers for α-globin genes are KU232561, KU232562, KU232563, KU232564, KU232565, KU232566, respectively (Additional file1). The phylogeny study of α-globin genes and their respective proteins within the group of these six sturgeon species and between this group and other fishes in NCBI revealed the trend of evolution of α-globin over the course of time.

The phylogenetic tree based on the α-globin sequences of sturgeons (including those in the present study) was superimposed on the morphological tree for the first time. The results indicated that the divergence of α-globin gene happened with higher speed in Caspian Sea sturgeon species, contemporaneously with the geological events leading to the formation of Caspian Sea.

This adaptation most likely represents these species’ adaptation to the new environmental conditions with variation in oxygen availability.

## Materials and methods

### Sequence analysis of the α-globin chain for six Caspian Sea sturgeon species

As there was no sequence information available for α-globin sequence of *Acipenser* in the literature, all fish α-globin amino acid sequences available at the NCBI database were retrieved (http://www.ncbi.nlm.nih.gov/protein/?term=alpha+globin+fish). Based on the alignment, a pair of degenerate intra-exonic primers Table 1 for PCR amplification of a portion of the gene were designed. The primers were used for amplification using genomic DNA obtained from *A.persicus, A.gueldenstaedtii, A.ruthenus, A.nudiventris, A.stellatus*, and *H.huso*. The genomic DNA isolated by conventional phenol-chloroform protocols [32] from red blood cell lysates of specimens obtained from the International Sturgeon Research Institute (Rasht, Iran).

**Table 1.**
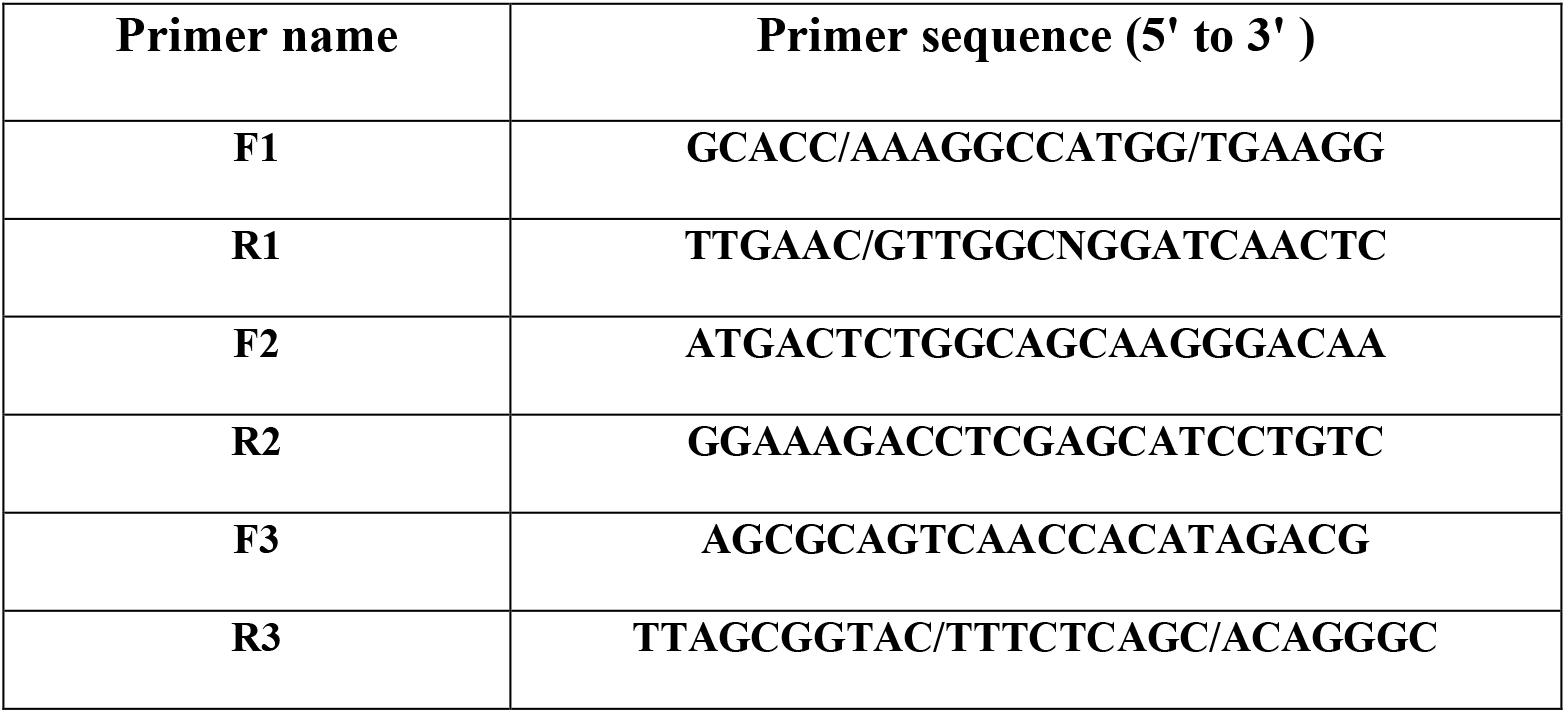
The pair of primers for PCR amplification of a portion of the gene were designed.

For the purpose of this study, no fish were killed. Five ml blood samples were collected in a non-invasive procedure from the caudal vein of alive fish as part of standard care. Each fish was out of the water for blood sampling between two to three minutes.

Animal procedures were approved by the Ethics Committee of University of Tehran under the ethics code IR.UT.REC.1395031.

Considering the reports [33] indicating the occurrence of polyploidization in some sturgeons and that our degenerated primers could amplify different alleles or paralogs of α-globin we used the following strategy criteria to amplify and compare unique sequences in our study. Only the distinguishable amplified bands were cut from the gel after electrophoresis of PCR products and sequenced after purification. In this way, it is less likely that paralogues sequences or other members of gene family with different size are mixed. Afterward, only the sequences with no ambiguity in their chromatogram were selected. Therefore it is less likely that alleles or similar sequences are combined. In the end, several primer pairs (Table 1) amplified overlapping regions of α-globin were used for sequencing α-globin gene, only sequences with perfect matches in their overlapping regions were selected.

The amplified bands were sequenced by the Sanger dideoxynucleotide protocol, high quality sequences were submitted to NCBI. The GenBank accession numbers for sturgeon α-globin genes are listed in the Additional file1. We used sturgeon alpha-globins sequenced here and the -globins recovered from NCBI in further analyses.

### Phylogenetic analysis

Nucleotide and amino acid sequence alignments and generation of Neighbour-joining (NJ) trees [34] were performed using MEGA 6.0 [35]. As an agglomerative clustering method, the Neighbour Joining (NJ) method assumes unequal rates of mutation along each branch and produces tree with branch lengths proportional to estimated divergence along each branch ant in units of nucleotide substitutions per site. The NJ tree approximates the minimum evolution tree (that whose total branch length is minimum).

The transition-transversion 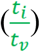 ratios are calculated using MEGA 6.0 and with Kimura 2-parameter (K2P) distances using both transitions and transversions [36].

The non-redundant and complete amino acid sequences of 110 fish α-globins from NCBI were selected.

The NJ analysis was performed on these sequences for tree construction in Fig.2. Then, for the construction of the phylogenetic tree in Fig. 3, we chose the exonic nucleotide sequences of 28 from 110 fishes belonging to different super-classes..

The phylogenetic analyses were carried out on the set of α-globin nucleotide sequences, including the 28 above mentioned sequences, in addition to the nucleotide sequences for α-globin genes of six sturgeons from this study. Phylogenetic trees were inferred by the NJ method. The robustness of all NJ trees was assessed by bootstrap analysis with 1000 replications.

### Divergence and Timetree analysis

There are various tools for inferring molecular timetrees and modeling rate variations among lineages such as PAML [37] and MULTIDIVTIME [38], Beast [39], DPPDiv [40], MCMCTree [41], MrBayes [42], PhyloBayes [43], r8s [44], and TreePL [45] that use Bayesian relaxed-clock dating as one of the most widely molecular dating techniques. We used the RelTime method [46] which estimates relative times of divergences for all branching points in phylogenetic trees based on the Maximum Likelihood method and without requiring specific knowledge for lineage rate distribution or specifying any clock calibrations and associated distributions. The RelTime has overcome many deficiencies in other mentioned methods. Estimating relative times implemented in MEGA 6.0 is useful for ordering the divergence time of α-globin genes [47].

For timetree calculation, we assume that the diversification of α-globin happened at the same time as morphological speciation.

Estimating of evolutionary divergence between exon nucleotide sequences, the ratio of transitional to transversional distances per site between 6 sturgeon α-globins sequences was estimated using MEGA 6.0 [35], see Table 2. Standard error estimates are shown above the diagonal. Analyses were performed using the K2P model [48]. The rate variation among sites was modeled with a gamma distribution (shape parameter = 1). All codon positions were used.. All positions containing gaps and missing data were eliminated. There was a total of 432 positions in the final dataset.

**Table 2.** Distance Matrix for sixα-globin Gene Sequences of Caspian Sea Sturgeons.

| *Sturgeon species* | **1** | **2** | **3** | **4** | **5** | **6** |
| --- | --- | --- | --- | --- | --- | --- |
| *1.A.persicus* |  | 0.447 | 0.575 | 0.689 | 0.742 | 0.442 |
| *2.A.gueldenstaedtii* | 0.331 |  | 0.377 | 0.416 | 0.416 | 0.317 |
| *3.A.ruthenus* | 1.559 | 0.913 |  | 0.344 | 0.419 | 0.338 |
| *4.A.nudiventris* | 1.538 | 0.874 | 0.819 |  | 0.441 | 0.307 |
| *5.A.stellatus* | 1.804 | 0.874 | 1.029 | 0.500 |  | 0.476 |
| *6.H.huso* | 1.165 | 0.742 | 0.952 | 0.605 | 1.022 |  |

## Results

### Phylogenetic analysis based on α-globin sequences is in accordance with the taxonomic position of the Caspian Sea sturgeons

Maximum likelihood (ML) and neighbor-joining (NJ) analyses of the nucleotide and amino acid sequences were performed to better ascertain the relation between the sequences of the various sturgeon species (Fig. 1).The trees inferred by both methods have similar structures.

**Fig. 1.**
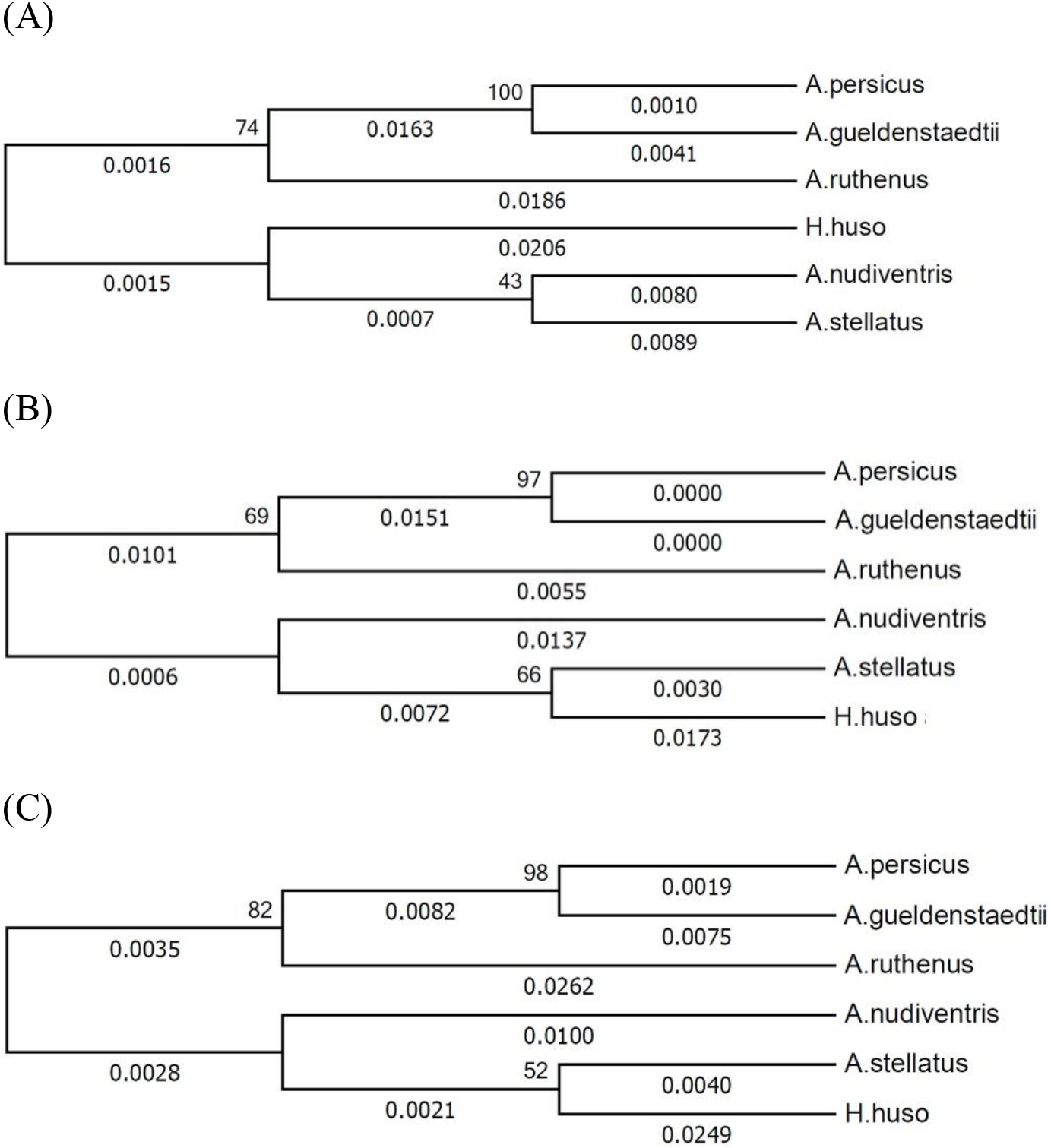
Phylogenetic tree based on α-globin estimated by the NJ algorithm. The branch lengths are proportional with number of base substitutions per site. The bootstrap percentages for 1000 replicates are shown next to the branches. Trees were inferred based on: (A) nucleotide sequences, (B) amino acid sequences, and (C) EST sequences.

The optimal NJ phylogenetic trees were inferred to determine the phylogenetic position of α-globin genes and proteins of 6 Caspian Sea sturgeons. Fig.1. shows the NJ phylogenetic tree based on (A) nucleotide sequences, (B) amino acid sequences, (C) EST sequences. The tree is drawn to scale, with branch lengths in the same units as the evolutionary distances used to infer the phylogenetic tree. The evolutionary distances were computed using the Maximum Composite Likelihood method and are in the units of the number of base substitutions per site. The percentage of replicate trees in which the associated taxa clustered together in the bootstrap test (1000 replicates) are shown next to the branches.

Figs 2 and 3 show the phylogeny position of sturgeon α-globins in a bigger picture composed of the representatives from all fish classes based on amino acid and EST sequences, respectively. In Fig.2, the Actinopterygii class, which includes sturgeons, has the most representatives in the tree and to avoid the unnecessary details, the position of globins from Actinopterygii class.

**Fig. 2.**
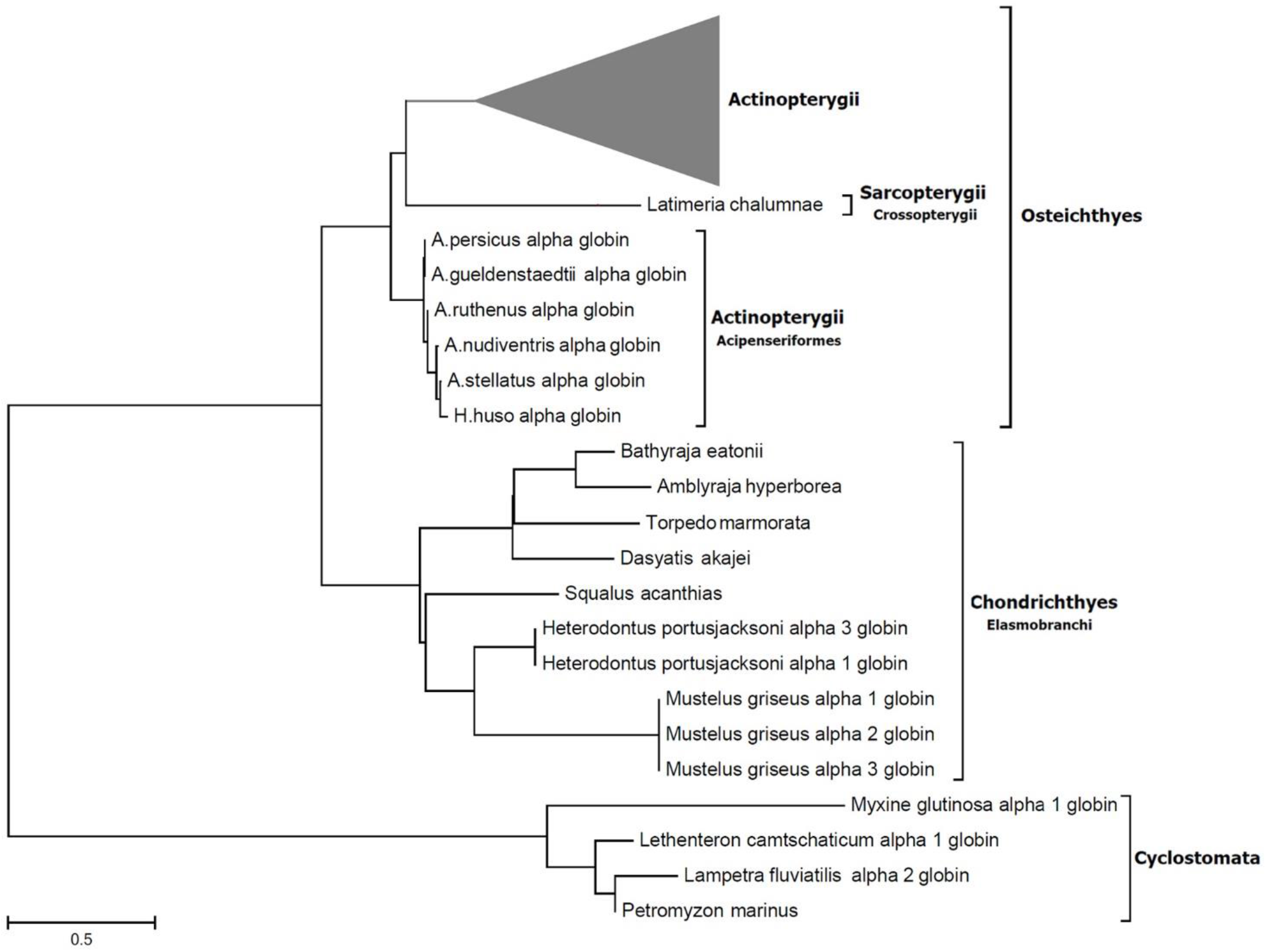
The NJ tree is drawn based on the 110 amino acid sequences of fish α-globins available at NCBI plus the six sturgeon species from current study. The triangle is the representative of α-globin proteins from Actinopterygii class.

**Fig. 3.**
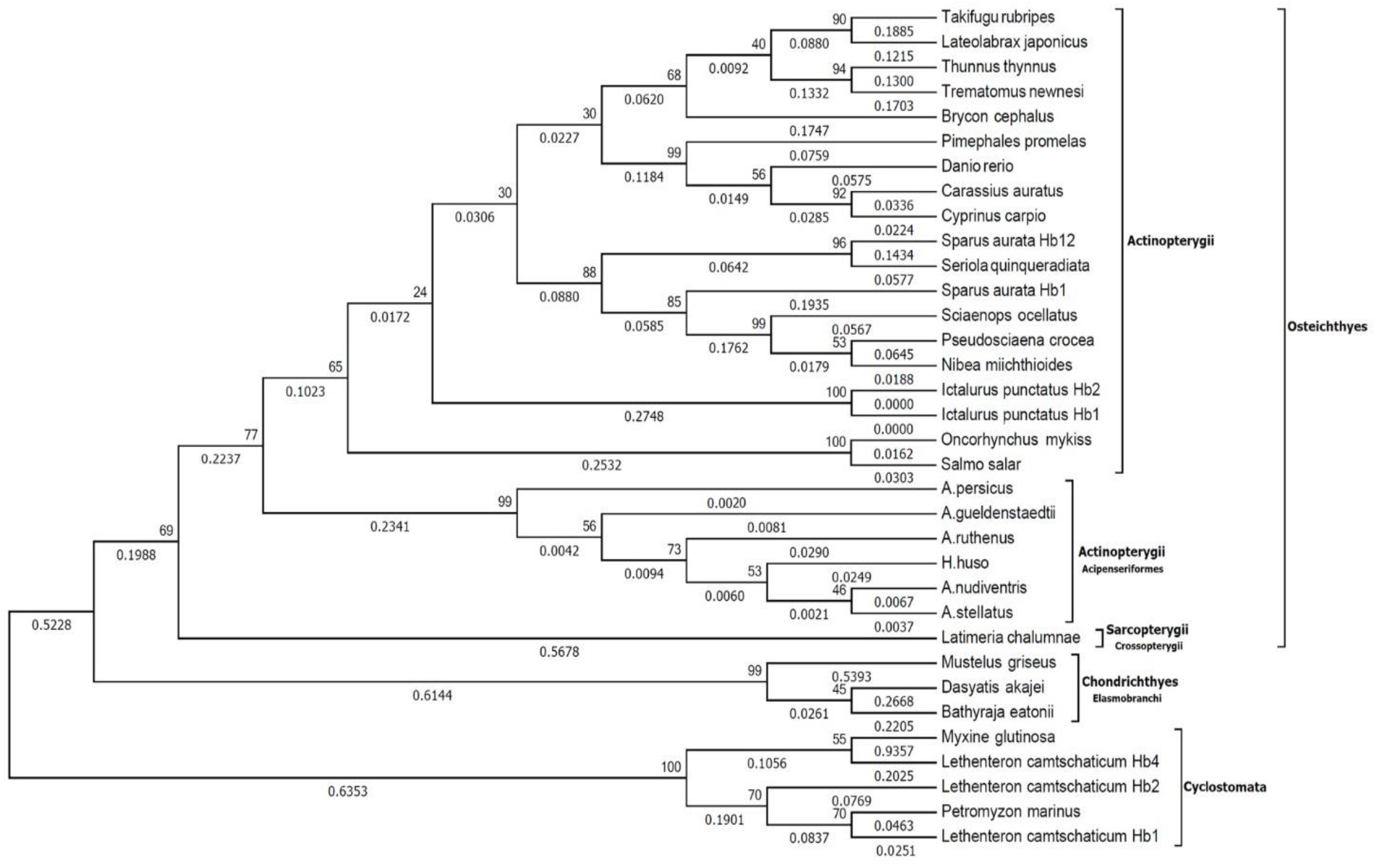
Phylogenetic NJ tree based on EST sequences of α-globin genes from 28 fish species plus six sturgeons in the current study. EST sequences properly discriminate between Chondrichthyes class, and Osteichthyes and Cyclostomata superclasses. The branch lengths are proportional with number of base substitutions per site. The bootstrap percentages for 1000 replicates are shown next to the branches. The capital letters refers to the corresponding clade in Fig. 5.

Figs 2 and 3 clearly show that both the α-globin amino acid and EST sequences could properly discriminate between Chondrichthyes class and Osteichthyes and Cyclostomata superclasses. This hierarchy is in accordance with the systematic classification of fishes.

For determining the phylogenetic position of the sturgeons based on the α-globin sequence, the 110 known α-globin chain amino acid sequences of fishes in NCBI, as well as the sturgeon sequences identified in this study, were compared by neighbor-joining analysis (Fig. 2).

The phylogenetic position of the Caspian Sea sturgeons was determined among other fishes. The tree was built by Neighbor-joining method based on α-globin chain amino acid sequences of the 110 fishes available at NCBI and the six sturgeon species of this study.

From the 110 α-globin amino acid sequences, the corresponding nucleotide sequences were extracted for 28 fish sequences presently available at NCBI. The phylogenetic analyses were carried out on the exon part of the set of α-globin nucleotide sequences, including the 28 above mentioned sequences and the nucleotide sequences for α-globin genes of six sturgeons from the current study. The phylogenetic tree was inferred by the NJ method assessed by bootstrap analysis with 1000 replications (Fig. 3).

The phylogenetic tree based on α-globin EST sequences from 28+6 fish species was created using the NJ method. Numbers on joints indicate the bootstrap values for 1000 replications and the numbers on branches show the measured genetic distances. The capital letters are referred to in Fig. 5 and intended to establish a relationship between molecular and morphologic phylogeny.

### Divergence of α-globin sequences occurred in a relatively short evolutionary period

The ratio of transitional to transversional distances per site between 6 sturgeon α-globins sequences was calculated to estimate evolutionary divergence between exon nucleotide sequences. The average evolutionary divergence over all sequence pairs based on this model was 0.982.

Distances were calculated with the program MEGA 6.0 using the Kimura two-parameter model [48] and the pairwise deletion option. The lower section shows the ratio of transitional to transversional distances per site between α-globin sequences of 6 sturgeon species calculated with a distribution to model different substitution rates between sites (shape parameter of the distribution, = 1). The upper section demonstrates standard error estimates.

The 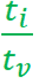 ratios can convey information about how recently separation of globins within the Acipenseriformes occurred. Values for 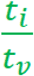 the approximately range between 0.331 and 1.804 for comparison between Acipenseriform species with an average of 0.982 and a variance of 0.40.

For comparative study, we excluded sturgeons of the Actinopterygii sub-class and then calculated the 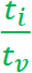 between the remaining members as one group and the six sturgeons as another group. The average 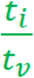 was 0.773.

In the current study, the RelTime algorithm was applied for determining the divergence time of α-globin nucleotide sequences. It also allows to assign age constraints to the corresponding nodes. To apply age constraints, 10 pairs of genera and species were selected from the phylogenetic tree in Fig.3 inferred by the neighbor-joining (NJ) method with 1000 bootstrap replications. Table 2 shows these pairs and their corresponding lower and upper borders and the median divergence time extracted from timetree official website [49,50].

Eleven pairs of families were selected to apply their divergence times as age constraints to calculate the divergence time of α-globin genes. The table contains lower and upper bounds of divergence times and the median extracted from the timetree official website.

A combined evidence approach (molecules plus geology) was adopted for assessing the relationship among the species included in Fig. 3. Fig.4 shows the timetree inferred from the Bayesian analysis based on the neighbor-joining tree of Fig. 3.

The tree inferred from the RelTime algorithm implemented in MEGA 6.0 for α-globin gene sequences. Branches are shown to reflect the estimated divergence times. The horizontal bar shows the divergence time in millions of years for the α-globin splits.

## Discussion

Earlier studies revealed some differences between the structure and function of Hbs in the Caspian Acipenseriformes clade [17,20,21].

In Fig.1, phylogenetic relationships are investigated for six Caspian Sea sturgeons. *A.persicus* and *A. gueldenstaedtii* α-globin sequences were consistently most similar (Additional file 2 and Additional file 3). This was expected because these species are considered closest to each other based on morphological taxonomy.

*Acipenser* and *Huso* are two separate genera ; however, *H.huso* falls in the *Acipenser* group based on molecular data. This is in accordance with our findings based on α-globin sequences as shown in Figures 1–3 (Additional file 2 and Additional file 3).

The minimum substitution of amino acids occurred between *A.persicus* and *A.guldenstaedti* (only one substitution), while maximal differences were observed between the sequences of *H.huso* and the other Acipenser species (Additional file 2 and Additional file 3).

*A.nudiventris* and *A.stellatus* nucleotide sequences were grouped together linked to *H.huso* in a subtree(Fig.1, (A)). However, at protein and EST levels, the positions of *H.huso* and *A.nudiventris* are exchanged (Fig.1, (B) and (C)). The high similarity between intron regions in *A.nudiventris* and *A.stellatus* (Additional file3) is the reason for this change in subtree topology.

From a holistic perspective, trees in parts (B) and (C) of Fig.1 are completely identical. This leads to conclude that exon nucleotide substitutions in the Caspian Sea sturgeons have not been deep enough to change the phylogeny positions of protein sequences.

A complete fossil record would allow for an extensive understanding of the evolutionary history of the group. However, the fossil record of fishes is rather incomplete.

The fact that a significant number of fish globin sequences is available in NCBI; provides adequate material to study the functional aspects of globin chains and their phylogenetic relationship.

The trees in Figs. 2 and 3 demonstrate the phylogenetic position of α-globins for fishes of different classes concerning protein and nucleotide sequences. Like for sturgeons, adaptation is evident for many teleost fishes, a group very near Acipenseriformes in the phylogenetic hierarchy (Fig.2). For example, *Anopiopoma fimbria, Gadus morthua, Osmerus mordax, Onncorthynchus mykiss, and Salmo salar* have a migratory lifestyle. These fish can tolerate a wide range of environmental conditions between the sea and freshwater. Various biological adaptations allow survival under variable conditions encountered in these two habitats, and features of Hb chains may be among the adaptations. Another example would be *Latimeria chalumnae* which is positioned near the Acipenseriformes in molecular phylogenetic trees [51]. There was a missing link between teleosts and Acipenseriformes in morphologically based phylogenetic trees. *Latimeria chalumnae*, a living fossil, was found in 1938 and filled the gap in the fossil record as it displayed little morphological differences compared to the Acipenseriformes [52]. Our phylogenetic tree based on α-globin sequences (Fig. 2) and its accordance with the morphological tree (Fig. 5) support the hypothesis that changes in Hb sequences have been part of adaptation processes. The summarized cladograms in the Fig.5 demonstrate the evolution on a morphologic basis [53–57] based on two sources of information. First, the knowledge acquired from modern ichthyology based on cladistic systematics and second, the timetree of life showing both branching order and times of divergence [50,58].

In this study, we used the RelTime algorithm [46,47] for inferring the evolutionary timetree. By default, the RelTime does not need to calibration points and produces estimates of relative times of divergence for all branching points in the phylogenetic tree without using clock calibrations. RelTime uses calibrations to translate relative times to absolute times. The relative times produced by the RelTime method can be directly converted into absolute times when a single known divergence time (calibration point) ia available based on fossil or other information [47]. In this study, for obtaining calibration points, we used the timetree official website [49,50,58,59]. Until today, the Timetree covers 3168 studies related to 97085 different species [http://www.timetree.org/references]. To apply age constraints, 10 pairs of genera and species were selected from the phylogenetic tree in Fig.3 inferred by the neighbor-joining (NJ) method with 1000 bootstrap replications. Table 3 shows these pairs and their corresponding lower and upper borders as well as the median divergence time extracted from timetree official website.

**Table 3.**
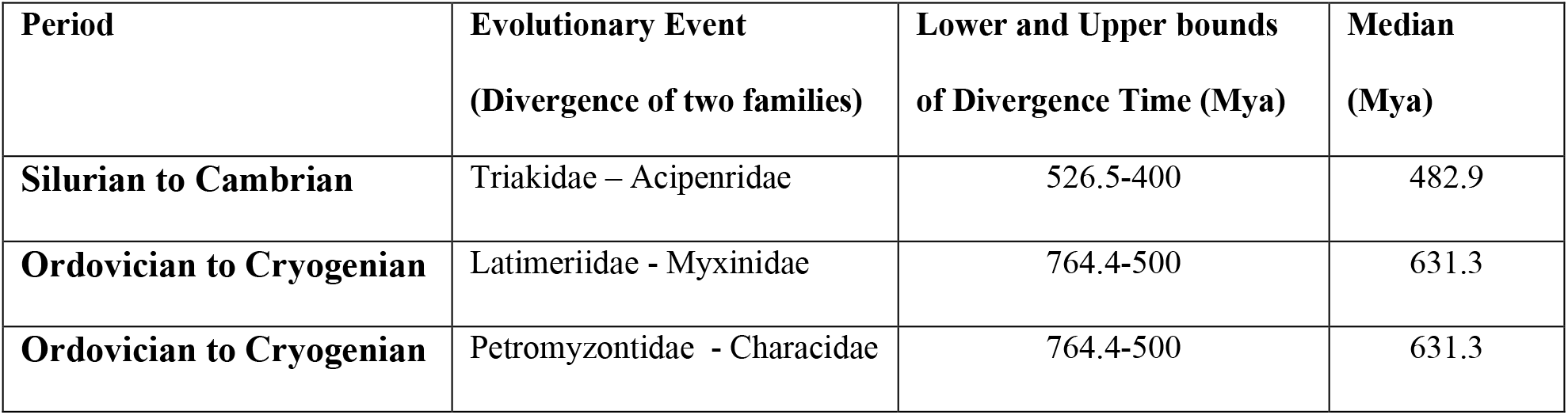
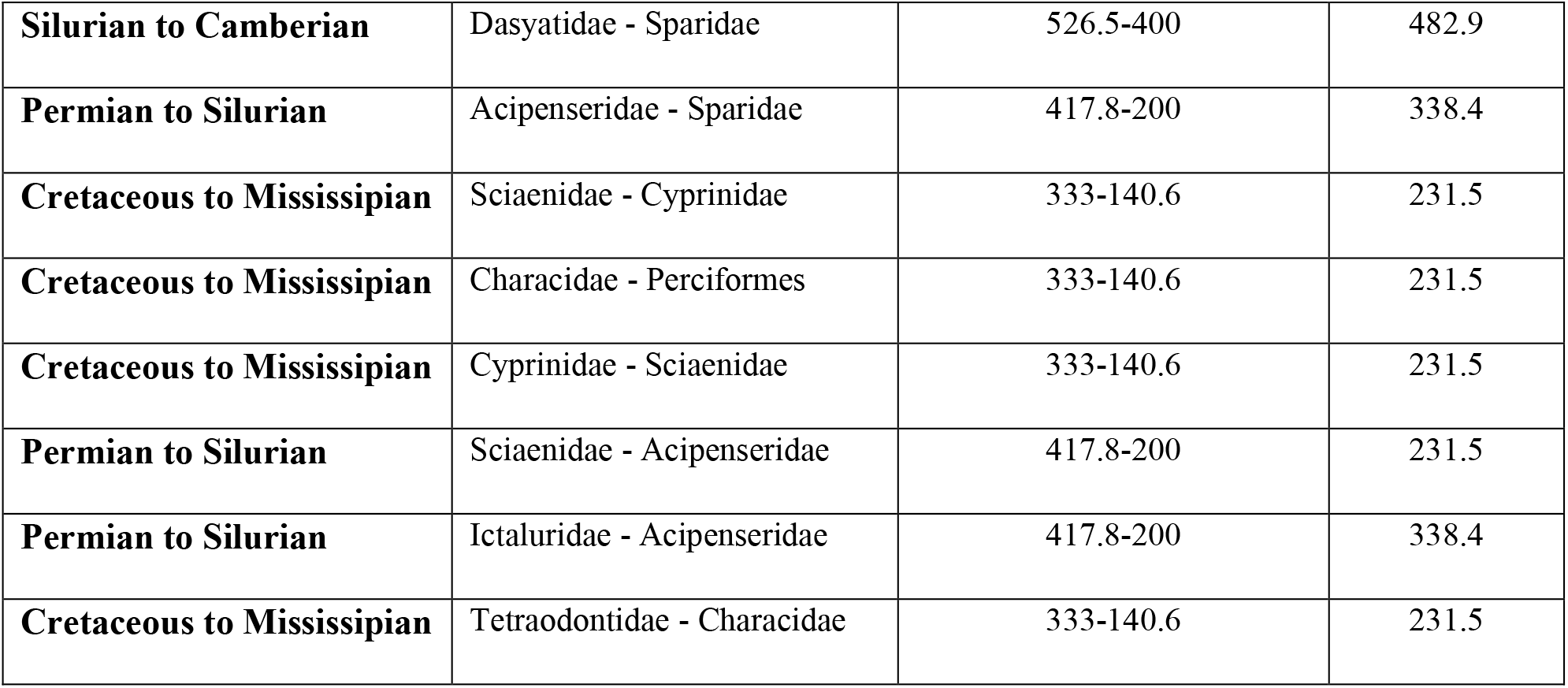
Geological Period and Time of Divergence for Different Fish Families.

| Period | Evolutionary Event<br>(Divergence of two families) | Lower and Upper bounds<br>of Divergence Time (Mya) | Median<br>(Mya) |
| --- | --- | --- | --- |
| **Silurian to Cambrian** | Triakidae – Acipenridae | 526.5-400 | 482.9 |
| **Ordovician to Cryogenian** | Latimeriidae - Myxinidae | 764.4-500 | 631.3 |
| **Ordovician to Cryogenian** | Petromyzontidae - Characidae | 764.4-500 | 631.3 |
| **Silurian to Camberian** | Dasyatidae - Sparidae | 526.5-400 | 482.9 |
| **Permian to Silurian** | Acipenseridae - Sparidae | 417.8-200 | 338.4 |
| **Cretaceous to Mississippian** | Sciaenidae - Cyprinidae | 333-140.6 | 231.5 |
| **Cretaceous to Mississippian** | Characidae - Perciformes | 333-140.6 | 231.5 |
| **Cretaceous to Mississippian** | Cyprinidae - Sciaenidae | 333-140.6 | 231.5 |
| **Permian to Silurian** | Sciaenidae - Acipenseridae | 417.8-200 | 231.5 |
| **Permian to Silurian** | Ictaluridae - Acipenseridae | 417.8-200 | 338.4 |
| **Cretaceous to Mississippian** | Tetraodontidae - Characidae | 333-140.6 | 231.5 |

The capital letters are intended to establish a one to one correspondence between the clades in this figure (based on morphology) and Fig. 3 (based on molecular phylogeny).

Comparing the sub-trees A to F in Figs.3 and 5, reveals a high level of similarity between trees and a one to one homology between labeled clades.

The comparison of sturgeon clade in Fig.3 with Chondrostei subclass in Fig.5 which includes Acipenseriformes order (Both marked with capital E), reveals that the α-globin conveys the potential information for phylogenetic clustering of fishes at least at the subclass level. But, because globin genes are not classified as stationary genes, it is not plausible to make the systematic phylogeny of organisms only based on them. A stationary gene, is considered as a gene which frequencies of its nucleotides do not change significantly among taxa [60]. Deviations from this stationary condition at any codon position can result in non-stationary conditions. Enough number genes should be exploited to infer a confident phylogenetic tree.

This data strongly support the hypothesis that the morphological speciation time of fishes in the course of evolution coincides with the α-globin divergence and both occur in the same geological period (Fig. 4). This simultaneity may emphasize the essential role of globin chains for survival of fishes under new environmental conditions reflecting the natural selection pressure that leads to speciation. The Caspian Sea, a remnant of the ancient ocean Paratethys Sea became landlocked in the recent Neogene geological period (2.58-23.03 Mya) due to tectonic uplift and a fall of sea level. Separation from the ocean, caused drastic changes in the physico-chemical conditions of the Caspian Sea, including salinity, temperature, and oxygen partial pressure. This relatively short time span had not been wide enough to make extensive speciation in sturgeon α-globins. The higher rate of transitions for sturgeons shows that the transversion as a driving force for speciation needs more time to exert its effect. In the timetree of Fig. 4, the divergence times of α-globin genes were determined by applying the time constraints presented in Table 2. The timetree was inferred based on the assumption that the appearance of the species and their corresponding α-globin proteins occurred simultaneously. Therefore, the validity of the timetree depends on the accuracy of this assumption.

**Fig. 4.**
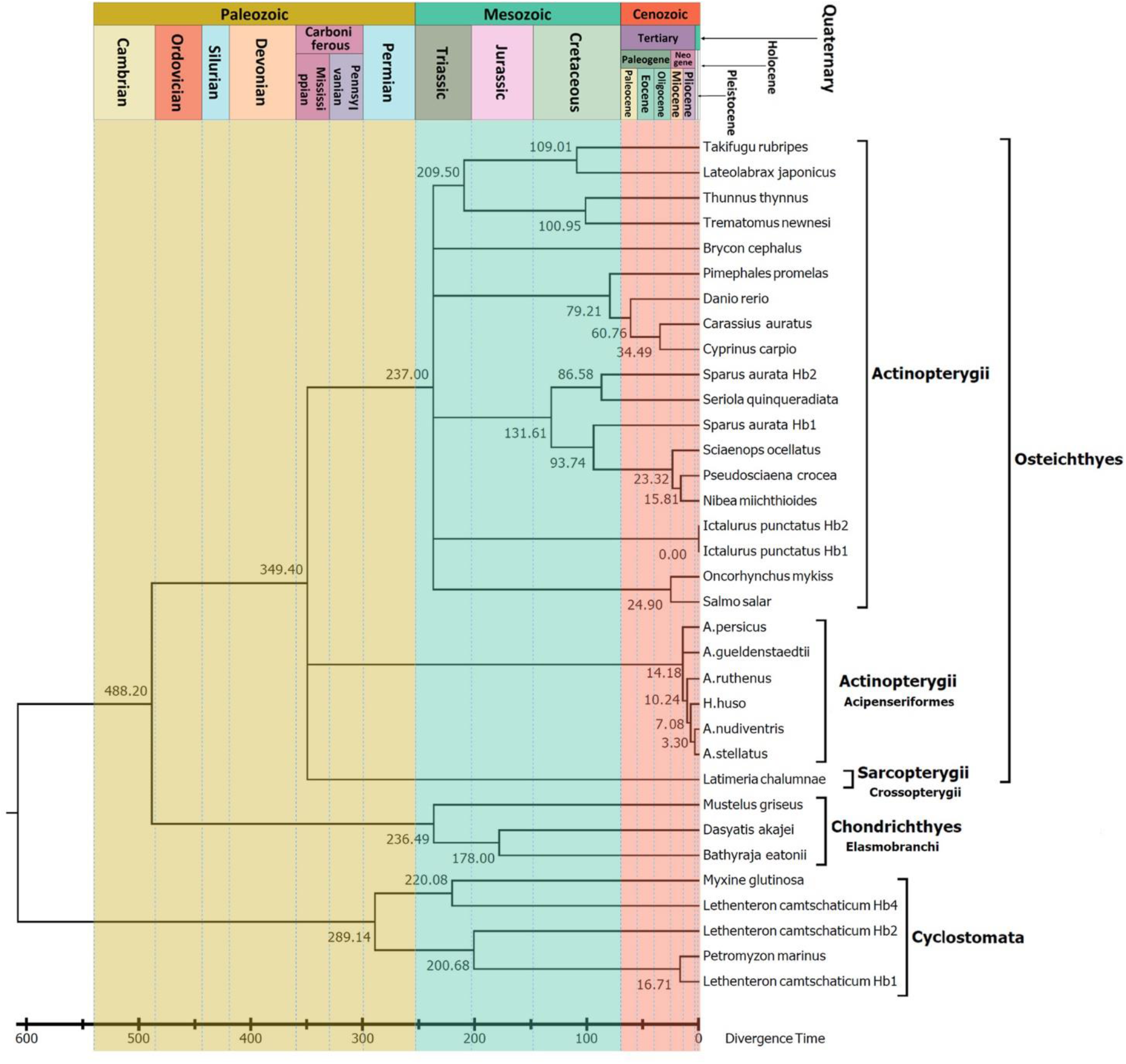
Rooted time tree for α-globin gene sequences inferred by the RelTime algorithm based on the NJ tree of Fig. 3. The branch lengths are proportional to the speciation time from the ancestor node. The absolute estimated speciation time is shown next to the branches in Mya.

**Fig. 5.**
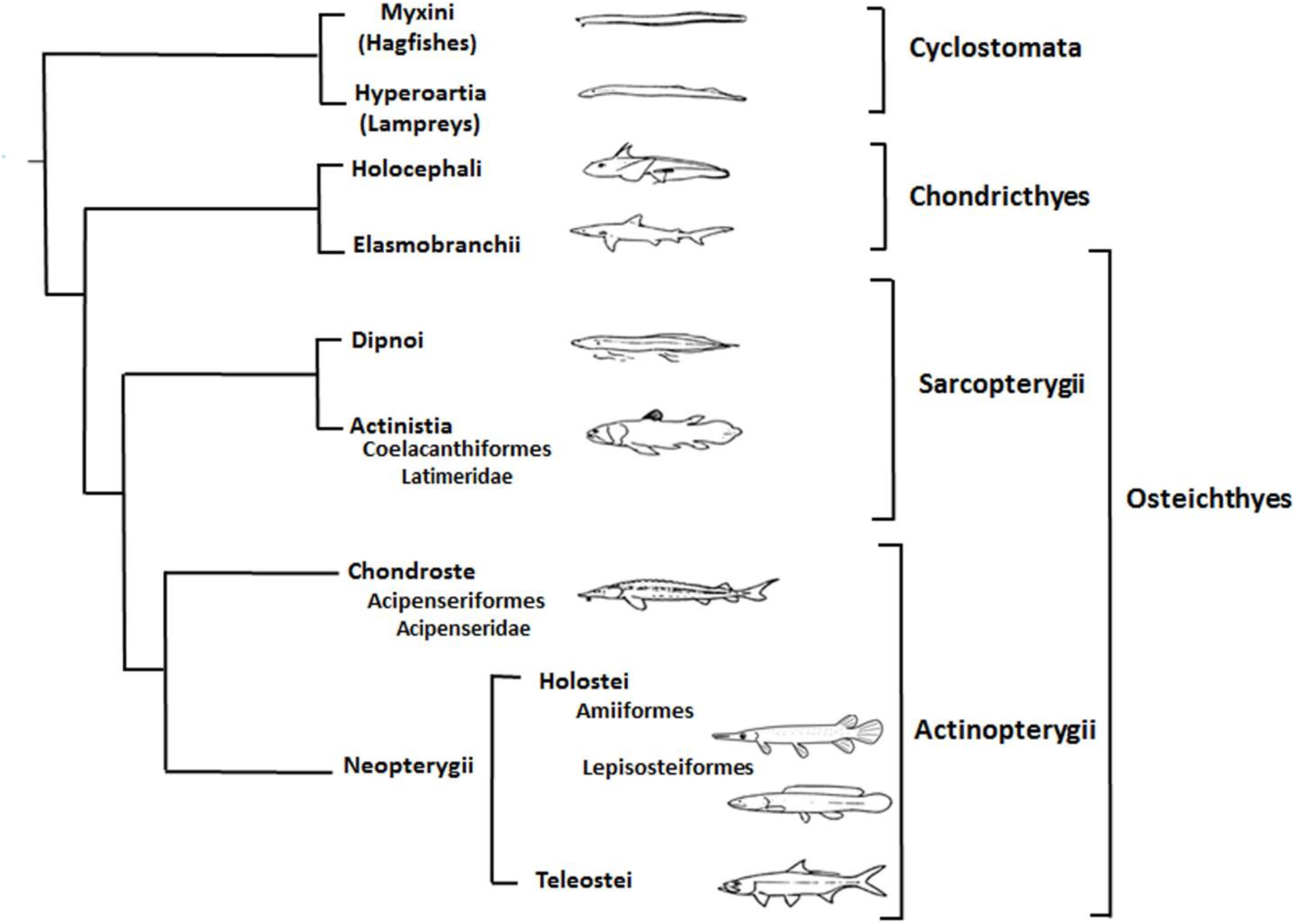
Cladogram showing the relationships of the Acipenseriformes order (capital letter E) to other fishes based on morphologic relationships. The capital letters are intended to show the correspondence between the positions of clades in morphologic phylogeny and clustering based on the α-globin genes illustrated in Fig. 3.

Among six sturgeons investigated in this study, only the *Acipenser persicus* is endemic of the Caspian Sea. According to Peng et al. (2007) [33], the divergence time of this species from its ancestor returns to approximately 10.8 Mya based on Cytochrome b which is completely in accordance with our analysis based on α-globin estimating the time 14.18 Mya. The only discrepancy for the dating belongs to *H. huso* (10Mya in our study compared to 80Mya in their study). Although the position of these species in the phylogenetic tree in both studies are similar (*A. Persicus* and *A. gueldenstaedtii* are grouped together while *H.huso* and *A. nudiventris* are closer to each other), the apparent differences in dating could be raised at least for two reasons. First the algorithm which is used in these two studies for dating of speciation are different. There are several methods for estimation of speciation time based on molecular clock concept. Some of these models, like MultidivTime which is used in Peng et al. (2007), are not sensitive to the ratio of 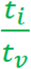 while some others like RelTime which is used in our study are sensitive to the 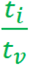. Calculation of 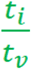 ratio for the sequences of cytochrome b for the fishes in Figure 4 of our study showed that indeed the cytochrome b of the Caspian Sea sturgeons have a higher ratio of 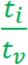 compared to their α-globin genes indicating α-globin gene have been under positive selection during the Caspian Sea formation. At the other hand the sampling of these two studies are different. While the collected sample of *H. huso* in our study, for instance, was from the Caspian Sea, in case of Peng et al. (2007) it was from other regions or GenBank. One possibility is that the α-globin gene of individuals in the Caspian Sea have different rate of substitution compared to the individuals in other geographical places where Peng et al. (2007) have collected their samples. Although this difference in α-globin sequences is the consequence of local adaptation, it has not lead to morphological changes. It is also tempting to speculate that in these two studies for these particular species where discrepancies observed in dating, by chance different orthologues of α-globin are analyzed as the *H.huso* and *A. nudiventris* are diploid while *A. persicus* and *A. gueldenstaedtii* are tetraploid [33].

The 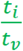 ratios can be used to suggest how recently separation of α-globins within the Caspian Sea Acipenseriformes occurred. Considering the long evolutionary history of *Acipenser*, it is expected that the transition substitutions have been saturated and therefore the average value of 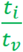 is reduced.

The average value of 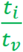 within Acipenseriformes α-globin is 0.98 and this value for comparisons between the Actinopterygii class and Acipenseriformes (excluding Sturgeons) is approximately 0.77. The higher average value within Acipenseriformes indicates that transition nucleotide substitutions are relatively less saturated than the sturgeons group.This is in contrast with the observations for the other fishes and can be best explained by more recent geological and climatic changes in the Caspian Sea.

## Conclusions

Sturgeons are living fossils and among the oldest surviving vertebrate species. Caspian Sea sturgeons are anadromous, which means they are able to cope with variable habitat conditions, a trait that demanded specific adaptations of oxygen-carrying hemoglobin molecules during evolutionary diversification. We ascertained the complete gene and protein sequences of an alpha-globin chain from six Caspian Sea sturgeons of different species. The amino acid and nucleotide sequences of α-globin genes were compared to all available fish α-globin sequences to reveal phylogenetic relationships and the divergence times. The fusion of information extracted from phylogenetic analysis of globin chains can throw light on the essentiality of proper changes in globin chains in the speciation process. This adaptation most likely represents the adaptation of these species to the new environmental conditions with variation in oxygen availability.

## Declarations

### Ethics approval and consent to participate

Not applicable

### Consent for publication

Not applicable

### Availability of data and material

The datasets generated and analyzed during the current study are available in the NCBI repository,

[https://www.ncbi.nlm.nih.gov/protein/1042177897, https://www.ncbi.nlm.nih.gov/protein/ANP93614.1,

https://www.ncbi.nlm.nih.gov/protein/ANP93613.1, https://www.ncbi.nlm.nih.gov/protein/ANP93612.1,

https://www.ncbi.nlm.nih.gov/protein/ANP93611.1, https://www.ncbi.nlm.nih.gov/protein/ANP93610.1].”

## Competing interests

The authers declare that they have no competing interests.

## Funding

The only funding for this research is limited to the materials and equipment which are provided by Institute of Biochemistry and Biophysics, University of Tehran and also Iran National Elites Foundation (INEF) and Iran National Science Foundation (INSF).

## Authors’ contributions

ShA and AMM delineated the rationale and developed the design of the study. ShA and KK engaged in sample collection. ShA and EE designed the experiments. ShA performed all experiments. ShA, KK, and AB analyzed the results and made conclusions.

All authors contributed to the final version of the manuscript and approved its content.

## Acknowledgements

The authors thank Dr. Ingo Schubert, Leibniz-Institute of Plant Genetics and Crop Plant Research (IPK), for improving the manuscript and fruitful discussion. We also thank the International Sturgeon Research Institute, Rasht, Iran, for providing samples and Mr. M. H. Tolouee for assistance.

## Authors’ information

Shohreh Ariaeenejad: Institute of Biochemistry and Biophysics (IBB), University of Tehran, Tehran, Iran-Department of Systems Biology, Agricultural Biotechnology Research Institute of Iran (ABRII), Agricultural Research Education and Extension Organization (AREO), Karaj, Iran

Kaveh Kavousi: Institute of Biochemistry and Biophysics (IBB), University of Tehran, Tehran, Iran Elaheh Elahi: School of Biology, College of Science, University of Tehran, Tehran, Iran

Ali Mohammad Banaei-Moghaddam: Institute of Biochemistry and Biophysics (IBB), University of Tehran, Tehran, Iran

Ali A. Moosavi-Movahedi: Institute of Biochemistry and Biophysics (IBB), University of Tehran, Tehran, Iran

## Notes

### Competing Interest Statement

The authors have declared no competing interest.

