## Additional file1 Table ,Additional file2 Table, Additional file3 Table for "Sturgeons: An Evolutionary Insight to The α-globin Protein Speciation and Diversification"

**Content**

Additional Tables S1 – S3

**Additional file1 Table. Accession numbers of α-globin sequences of the sturgeon species submitted to GenBank.** Each row shows species name, its Genbank accession number, and the length of submitted sequence.

**Additional file2 Table. Comparison of exon and intron parts of α-globin genes.** Two by two comparison of exon, intron, and total nucleotide sequences of α-globin genes among the sturgeon species. The numbers show the total nucleotide substitutions for two species among the same exons and introns. The last column shows total number of substitutions.

**Additional file3 Table. Comparison of amino acids of α-globin proteins.** Two by two comparison of total amino acid sequences of α-globin proteins among the sturgeon species. The numbers show the total amino acid substitutions for two species. The last column shows total number of substitutions.

**Additional file1 Table. Accession number of α-globin sequences the sturgeon species submitted to GenBank**

| Length of coding sequence (nucl.) | Accession number | *Sturgeon species* |
| --- | --- | --- |
| 812  812  901  868  875  825 | KU232561 KU232562 KU232563 KU232564 KU232565 KU232566 | *A.persicus*  *A.gueldenstaedtii*  *A.ruthenus*  *A.nudiventris*  *A. stellatus*  *H.huso* |

**Additional file2 Table. Comparison of exon and intron parts of α-globin genes.**

| Total differences | Exon 3 | Intron 2 | Exon 2 | Intron 1 | Exon 1 | *Sturgeon species* | |
| --- | --- | --- | --- | --- | --- | --- | --- |
| 4  126  78  94  92 | 0  1  0  2  7 | 0  2  3  4  27 | 3  12  7  5  6 | 0  109  65  79  48 | 1  2  3  4  4 | *A.gueldenstaedtii*  *A.ruthenus*  *A.nudiventris*  *A.stellatus*  *H.huso* | *A.persicus* |
| 4  128  81  96  94 | 0  1  0  2  7 | 0  2  3  4  27 | 3  15  10  8  9 | 0  109  65  79  48 | 1  1  3  3  3 | *A.persicus*  *A.ruthenus*  *A.nudiventris*  *A.stellatus*  *H.huso* | *A.gueldenstaedtii* |
| 125  128  78  58  122 | 0  1  1  1  6 | 2  2  2  3  27 | 12  15  13  11  13 | 109  109  59  39  72 | 2  1  3  4  4 | *A.persicus*  *A.gueldenstaedtii*  *A.nudiventris*  *A.stellatus*  *H.huso* | *A.ruthenus* |
| 78  82  78  32  68 | 0  0  1  2  7 | 3  3  2  3  27 | 7  10  13  2  5 | 65  65  59  23  25 | 3  4  3  2  4 | *A.persicus*  *A.gueldenstaedtii*  *A.ruthenus*  *A.stellatus*  *H.huso* | *A.nudiventris* |
| 94  96  58  32  81 | 2  2  1  2  5 | 4  4  3  3  28 | 5  8  11  2  3 | 79  79  39  23  41 | 4  3  4  2  4 | *A.persicus*  *A.gueldenstaedtii*  *A.ruthenus*  *A.nudiventris*  *H.huso* | *A.stellatus* |
| 92  94  122  68  81 | 7  7  6  7  5 | 27  27  27  27  28 | 6  9  13  5  3 | 48  48  72  25  41 | 4  3  4  4  4 | *A.persicus*  *A.gueldenstaedtii*  *A.ruthenus*  *A.nudiventris*  *A.stellatus* | *H.huso* |

**Additional file3 Table. Comparison of amino acids of α-globin proteins.**

| Total differences | Exon 3 | Exon 2 | Exon 1 | *Sturgeon species* | |
| --- | --- | --- | --- | --- | --- |
| 0  3  5  6  7 | 0  1  0  2  4 | 0  2  4  3  2 | 0  0  1  1  1 | *A.gueldenstaedtii*  *A.ruthenus*  *A.nudiventris*  *A. stellatus*  *H.huso* | *A.persicus* |
| 0  3  5  6  7 | 0  1  0  2  4 | 0  2  4  3  2 | 0  0  1  1  1 | *A.persicus*  *A.ruthenus*  *A.nudiventris*  *A.stellatus*  *H.huso* | *A.gueldenstaedtii* |
| 3  3  5  4  5 | 1  1  1  1  3 | 2  2  3  2  1 | 0  0  1  1  1 | *A persicus*  *A.gueldenstaedtii*  *A.nudiventris*  *A.stellatus*  *H.huso* | *A.ruthenus* |
| 5  5  5  3  6 | 0  0  1  2  4 | 4  4  3  1  2 | 1  1  1  0  0 | *A.persicus*  *A.gueldenstaedtii*  *A.ruthenus*  *A.stellatus*  *H.huso* | *A.nudiventris* |
| 6  6  4  3  3 | 2  2  1  2  2 | 3  3  2  1  1 | 1  1  1  0  0 | *A.persicus*  *A.gueldenstaedtii*  *A.ruthenus*  *A.nudiventris*  *H.huso* | *A.stellatus* |
| 7  7  5  6  3 | 4  4  3  4  2 | 2  2  1  2  1 | 1  1  1  0  0 | *A.persicus*  *A.gueldenstaedtii*  *A.ruthenus*  *A.nudiventris*  *A.stellatus* | *H.huso* |
